# Fine scale sampling reveals spatial heterogeneity of rhizosphere microbiome in young *Brachypodium* plants

**DOI:** 10.1101/2023.01.20.524947

**Authors:** Shwetha M. Acharya, Mon Oo Yee, Spencer Diamond, Peter F. Andeer, Nameera F. Baig, Omolara T. Aladesanmi, Trent R. Northen, Jillian F. Banfield, Romy Chakraborty

## Abstract

For a deeper and comprehensive understanding of the diversity, composition and function of rhizosphere microbiomes, we need to focus at the scale of individual roots in standardized growth containers. Root exudation patterns are known to vary across distinct parts of the root giving rise to spatially distinct microbial niches. To address this, we analyzed microbial community from two spatially distinct zones of the primary root (the tip vs. the base) in *Brachypodium distachyon*, grown in natural soil using standardized fabricated ecosystems known as EcoFABs as well as in more conventional pot and tubes. 16S rRNA based community analysis showed a stronger rhizosphere effect in the root base vs. bulk soil compared to the root tips vs. bulk soil, resulting in an enrichment of Actinobacteria, Bacteroidetes, Firmicutes and Proteobacteria, few OTUs belonging to less characterized lineages such as Verrucomicrobia and Acidobacteria. While the microbial community distributions are similar across growth containers, the EcoFAB displayed higher replicate reproducibility. Genome-resolved and bulk metagenomics revealed that genes associated with transcriptional regulation, transport of nutrients and catabolic enzymes indicating active metabolism, biofilm formation and root colonization were enriched in root tips. On the other hand, genes associated with nutrient-limitation and environmental stress were prominent in the bulk soil compared to the root tips, implying the presence of easily available, labile carbon and nutrients in the rhizosphere relative to bulk soil. Such insights into the relationships between root structure, exudation and microbial communities are critical for developing understanding of plant-microbe interactions.

## 1. Introduction

Plants exude 20-40% of their photosynthetically fixed carbon through intact root cells into the surrounding soil [1]. Besides root characteristics, root exudates are a key determinant for development of rhizosphere community. These root exudates containing low-molecular weight organic compounds, and together with mucilage and sloughed off root tissues mainly expelled from root tips, root exudates provide a major source of nutrients for the rhizosphere microbiome [2]. These compounds create a unique environment in the rhizosphere that is physiochemically distinct from the surrounding bulk soil and play a key role in recruiting and selecting relevant beneficial microbes to form a rhizosphere microbiome which is also distinctly differentiated from that of the surrounding bulk soil [3].

Root exudation patterns have been shown to vary spatially along the root system, exudates from rapidly dividing root tips differ in composition from exudates released from older sections of the root [4]. While the assembly of microbial community along different parts of roots (biogeography) is considered an important parameter in rhizosphere dynamics, systematic and standardized studies probing this deeper are lacking. Most rhizosphere microbiome studies, where plants are grown in soil, do not compartmentalize the roots based on their morphology but rather based on radial distance from the root axis (rhizosphere, rhizoplane and endosphere). As a result, capturing the effect of spatial differences along the roots is much unexplored, causing a gap in understanding how these differences impact microbial assembly in the rhizoplane.

Furthermore, while few studies in the past have demonstrated influence of plant growth container type on plant morphology [5–9], direct impacts of growth containers on the rhizosphere microbiome is relatively unexplored. Complex biochemical processes and interactions occur in microscale dimensions surrounding the root as outlined above. The ability to interrogate these processes within highly reproduceable and controlled growth containers will propel our understanding of rhizosphere spatial heterogeneity [10].

In this study, we investigated rhizosphere biogeography from two distinct root zones of *Brachypodium distachyon* grown in natural soil but in three different types of growth containers-conventional pots, tubes as well as specially fabricated EcoFABs [11] to assess (a) microbiome structure and function across root tips, root base and bulk soil; and (b) the suitability of standardized growth containers to study plant-microbe interactions at such finer scales. We also tested these different containers under open or closed environments (encased within secondary containment). The EcoFABs had demonstrated to be of high value in standardized investigations of plant traits and microbiome, and have been shown to reproducibly produce plant phenotypic traits and metabolite production [12], but their applicability to study spatially resolved rhizosphere had been hitherto unexplored. We used long read 16S rRNA amplicon sequencing and shotgun metagenomic sequencing to delineate differences between these diverse containers and distinct root zones (root tips, root base). Metagenomic functional potential unraveled significant differences between root tips and bulk soil.

## 2. Materials and Methods

### 2.1 Soil and plant growth conditions

Soil for plant growth was collected from the south meadow field site at the Angelo Coast Range Reserve in northern California (39° 44′ 21.4′′ N 123° 37′ 51.0′′ W) in August 2020. The upper layer (0-10 cm) was collected in clean collection bags, immediately transported on ice and stored at 4°C until further processing. The collected soil was passed through a 2 mm sieve to remove larger particles like dry roots and rocks prior to use.

In this study, we used three types of containers, EcoFAB, test tubes and plastic pots to grow *B. distachyon* (Bd21-3 plant line). EcoFABs (n = 11) were fabricated as reported earlier [13] with slight modifications. Briefly, the oval-shaped polydimethylsiloxane (PDMS) cast measuring 7.7 cm x 5.7 cm x 0.5 cm (height x width x depth) providing a container volume of 10 mL was held together by metal clamps and screws. Sterile plastic test tubes (n = 14) used to grow plants were 10 cm long with a diameter of 1.5 cm, and had a hole drilled at the bottom to drain excess water. The pots (n = 14) used were 10 cm x 10 cm squares with a depth of 10.5 cm, tapered from top to bottom. The volume of soil in test tube and EcoFAB was kept at 15 g each while the pot contained 600 g. The vertical distance between the sown seed to the bottom of the container was 8 cm for EcoFAB and 9 cm for both pot and test tube. Except for soil, all components were sterilized by UV sterilization or autoclaving. In addition, approximately half of all containers were kept sterile in closed Microbox containers (Sac O2, Belgium) while others were kept open to the environment.

Cold-treated *Brachypodium distachyon* seeds were de-husked, surface-sterilized in 70% ethanol followed by 50% household bleach for 5 minutes each and rinsed thoroughly in sterile water. They were germinated on sterile 0.8% noble agar plates under sunlight at room temperature for two days. Germinated seedlings were transferred into the containers taking care to place it 0.5 cm below the soil surface, watered once at 100% capacity with sterile water. Subsequent watering was done at 15% holding capacity, every 2 and 4 days for the open and closed containers respectively. The plants were placed in a greenhouse with a 16-hour photoperiod, 87.5% relative humidity, and average day and nighttime temperatures of 19.9 °C and 17.9 °C respectively.

### 2.2 Plant phenotypic measurements

Plants were harvested from all containers 14 days after sowing when the primary root had reached bottom of EcoFAB, and key plant phenotypic characteristics were measured. After excising the roots from the base of plant shoot, dry shoot weight was obtained by oven drying the shoots at 80 °C for 24 h followed by cooling to room temperature and measuring dry weight [14–16]. Shoot length was measured from end of the longest leaf to the point where root starts [17]. Root length was measured from root base to tip of the primary root.

### 2.3 Rhizosphere and bulk soil sample collection

At the time of harvest, roots were excised carefully from soil under aseptic conditions and lightly shaken to remove loosely attached bulk soil. Root tip and root base samples were harvested as 2 cm cuttings, measured from tip of the root, and from base of the plant shoot respectively. Due to complications during sampling resulting in physical damage to the roots, some samples were discarded reducing the number of root samples to n = 8, n = 11, and n = 7 originating from EcoFAB, test tube, and pot respectively. The loosely-bound rhizosphere soil was obtained by vortexing the root in 5 mM sodium pyrophosphate for 15 seconds, three times. The root was then placed in fresh pyrophosphate buffer and sonicated for 5 mins to extract tightly-bound fraction. To ensure the complete representation of the rhizosphere microbiome, both the loosely- and tightly-bound fractions were pooled for subsequent DNA extraction. Bulk soil (0.5 g) was collected from containers at least 1 cm away from the roots and kept frozen before DNA extraction.

### 2.4 DNA extraction and sequencing

Genomic DNA was extracted using DNeasy PowerLyzer Powersoil kit (Qiagen, US) following the manufacturer’s instructions and the eluted genomic DNA was quantified using Qubit™ dsDNA High Sensitivity assay kit (Thermofisher, US).

For bacterial full-length 16S rRNA amplication and sequencing, genomic DNA from all the available different root locations and bulk soil were sent to Loop Genomics (US). Briefly, the DNA was amplified with indexed forward (5’ CTGCCTAGAACA [Index, F] AGAGTTTGATCMTGGCTCAG 3’) and reverse primers (5’ TGCCTAGAACAG [Index, R] TACCTTGTTACGACTT 3’) and sequenced using the Illumina sequencing platform via paired end (150bp X 2) mode followed by the standard Loop Genomics informatics pipeline that uses short reads to construct synthetic long reads [18].

For metagenomic sequencing, replicates of each sample type (root tip, root base or bulk soil from each type of container) was pooled to accommodate the 200 ng DNA concentration requirement, resulting in a total of 9 samples. These samples were sent to QB3-Berkeley Functional Genomics Laboratory (University of California, Berkeley, US) (http://qb3.berkeley.edu/fgl/) for library prep and subsequent sequencing using Illumina 150 bp X 2 paired end reads with a depth of 20 Gb per sample.

### 2.5 16S rRNA community analysis

16S amplicon samples which contained less than 1000 reads after demultiplexing were discarded before analysis. We ensured that there were at least 3 replicate samples for every type of sample under the three variables tested; 1. Container (EcoFAB, pot or test tube), 2. Location (root tip, root base or bulk soil) and 3. Condition (Closed or Open). The demultiplexed data from loop genomics was then clustered into OTUs using usearch (version 11.0.667) for comparative analyses as follows [19]. Briefly, FASTQ files were 1st trimmed (1400 bps) and quality filtered (maximum expected error cutoff 1.0) before initial clustering and chimera filtering using Unoise 3 command. The resulting OTUs were further clustered to 97% identity before generating the OTU table, taxonomic assignments and comparative analyses.

From the OTUs generated through usearch, DECIPHER v2.0 (r studio package) was used to obtain taxonomic information based on the SILVA SSU version 138 [20, 21] following default parameters. The generated OTU samples were subjected to Hellinger transformation using decostand method in vegan R package version 2.5-7 [22] to standardize differences in sequencing depth prior to diversity analysis. Differential abundance of microbial OTUs across different containers and sample locations were determined using the DESeq2 package (version 1.14.1) in R [23]. Pairwise comparison between sample locations coupled to each container was carried out using a full DESeq2 model (design = ∼Container_Location + Condition). OTUs showing significant log-fold changes (p_adj_ <0.05) in at least one of these comparisons was further selected and visualized on a phylogenetic tree in iToL [24]. The log fold-change values were tested for correlation using Spearman’s test through custom python script. Afterwards, pairwise comparisons were repeated with a reduced model (design = ∼Location + Container + Condition) to study the effect on sample location while controlling container and condition variations. Using the transformed data, homogeneity of multivariate dispersions was analyzed for each sample location in each container using betadisper from vegan R package.

### 2.6 Metagenome assembly, annotation, and binning

Shotgun metagenomic sequence for the 9 samples (3 containers * 3 locations) were individually assembled using IDBA-UD v1.1.3 [25] with the parameters: -pre_correction -mink 20 -maxk 150 -step 10. Following metagenome assembly, all samples were filtered to remove contigs smaller than 1 kb using pullseq (https://github.com/bcthomas/pullseq). Open reading frames were then predicted on all contigs using Prodigal v2.6.3 [26] with the parameters: -m -p meta. KEGG KO annotations were predicted using KofamScan [27] using HMM models from release r02_18_2020 using default options. In cases where multiple HMMs matched a protein above threshold, the HMM with the lowest E-value had its annotation transferred to the protein.

Metagenome assemblies were binned into draft genomes using a combination of 4 automated binning methods. Briefly, reads from all 9 samples were mapped to assembled contigs ≥ 2.5□kbp using Bowtie2, and a differential coverage profile for each contig across all samples was used as input for the following differential coverage binners: MaxBin2, CONCOCT, vamb, and MetaBAT [28–31]. The algorithm DasTool [32], was then used to select the highest quality bins across the 4 binning outputs for each metagenome assembly. Finally, the full genome set across all samples (n = 146 genomes) was de-replicated at the species level (Average Nucleotide Identity ≥ 95%) using dRep [33] with the following parameters: -p 16 -comp 10 -ms 10000 -sa 0.95, resulting in a total of 42 species representative genomes. Species representatives were further selected to have ≥ 60% completeness and ≤ 10% contamination as estimated by checkM [34], this resulted in a final set of 32 species representative genomes meeting the criteria. 16S rRNA sequences were extracted from genomes with ContEst16S tool available online (https://www.ezbiocloud.net/tools/contest16s, last accessed on August 17, 2022) [35]. These 16S rRNA sequences were compared with the OTUs obtained from amplicon sequencing using BLAST+ [36] to check for taxonomic consistency.

### 2.7 Phylogenetic and abundance analysis of genome bins

Phylum level taxonomic assignments of 32 de-replicated genome bins and 1 genome (*P*. calidifontis - GCA000015805) included as an outgroup were inferred using GTDB-Tk v1.5.1 [37] with reference data version r202; phylogenetic relationships between de-replicated genome bins were inferred using GToTree v1.5.22 based on a set of 25 marker genes, and a phylogenetic tree was produced using FastTree2 [38].The tree was displayed and rooted in Geneious Prime v2020.2.4. The relative abundance of the 32 genome bins in all samples was assessed by cross mapping reads from each of the 9 samples back to the genome bins using Bowtie2, followed by quantification of coverage of genomes in each sample using coverM (https://github.com/wwood/CoverM). Differential abundance of genomes between rhizosphere spatial locations was assessed using the DESeq2 package in R [23]. Detailed version of this section can be found in Supplementary material.

### 2.8 Bulk Metagenome Analysis

Phylum level taxonomic composition of bulk metagenomes was assessed directly from raw sample reads using graftM [39] run with a custom ribosomal protein L6 (rpL6) marker database constructed from the r202 release of the GTDB database. Differentially abundant KO genes across the different sample locations were determined using the DESeq2 package (version 1.14.1) in R [23]. Pairwise comparison between sample locations was carried out using a reduced DESeq2 model (design = ∼Location). Heatmap of differentially abundant genes were plotted in R using the variance stabilized abundance values.

## 3. Results

### 3.1 Container type has minimal impact on plant phenotypic growth

We investigated the spatial biogeography of rhizosphere microbiome of *B. distachyon* grown in model fabricated ecosystems (EcoFABs) in comparison with conventional containers. *B. distachyon*, a model grass species for wheat family, was chosen as it produces only one fine primary axile root from the base of the embryo [40] on which the microbial spatial analysis was performed.

We measured three major phenotypes of plant growth, i.e., dry shoot weight, shoot length, root length, to determine container impacts on general plant growth. The only significant difference was between plants grown in pots in open vs. closed conditions (**Fig. S1**). The microbox used to maintain sterile condition (closed) was observed to trap a visibly higher amount of moisture inside the box and likely created higher water retention promoting plant growth. Regardless, no other significant difference was detected within or among containers despite differences in container architecture.

### 3.2 Location on root is the highest driver of microbial community dissimilarity

We analyzed the rhizosphere microbial community from two different root locations of a 14-day old *B. distachyon* and the bulk soil using full length 16S rRNA obtained using synthetic long read technology. Among the 3674 OTUs obtained after quality filtering, 25 different phyla were identified which corresponded to approximately 80% - 87.5% of all reads among the samples. Microbial relative abundance showed on average a dominance of the bacterial phyla Proteobacteria (22.3%-29.3%), Actinobacteriota (14.2%-23.5%), Acidobacteriota(12.2%-16.5%), Chloroflexi(6.3%-10.1%), Planctomycetota (3.7%-4.7%), Verrucomicrobiota (4.2%-7.4%), Bacteriodota (1.6-4.5%) and Myxococcota (1.9-2.6%) in all samples (**Fig. 1a**). Interestingly, phyla Firmicutes had lower relative abundance in bulk (average -0.4%) compared to root tip and root base samples (average - 2.6%). Microbial diversity was lower in root tip compared to bulk soil (p<0.005, Anova and Tukey) or root base (p<0.05, Anova and Tukey) in all three alpha diversity metrics analyzed (species number, Shannon and inverse Simpson) (**Fig. S2**). On the other hand, no significant difference in diversity was observed between root base and bulk soil. When compared between the containers for each sample location, for instance, root tip samples between the three containers, there was no significant difference in microbial diversity (p>0.05, Anova and Tukey) indicating negligible container impact. The same was observed for root base and bulk soil sample locations.

**Figure 1.**
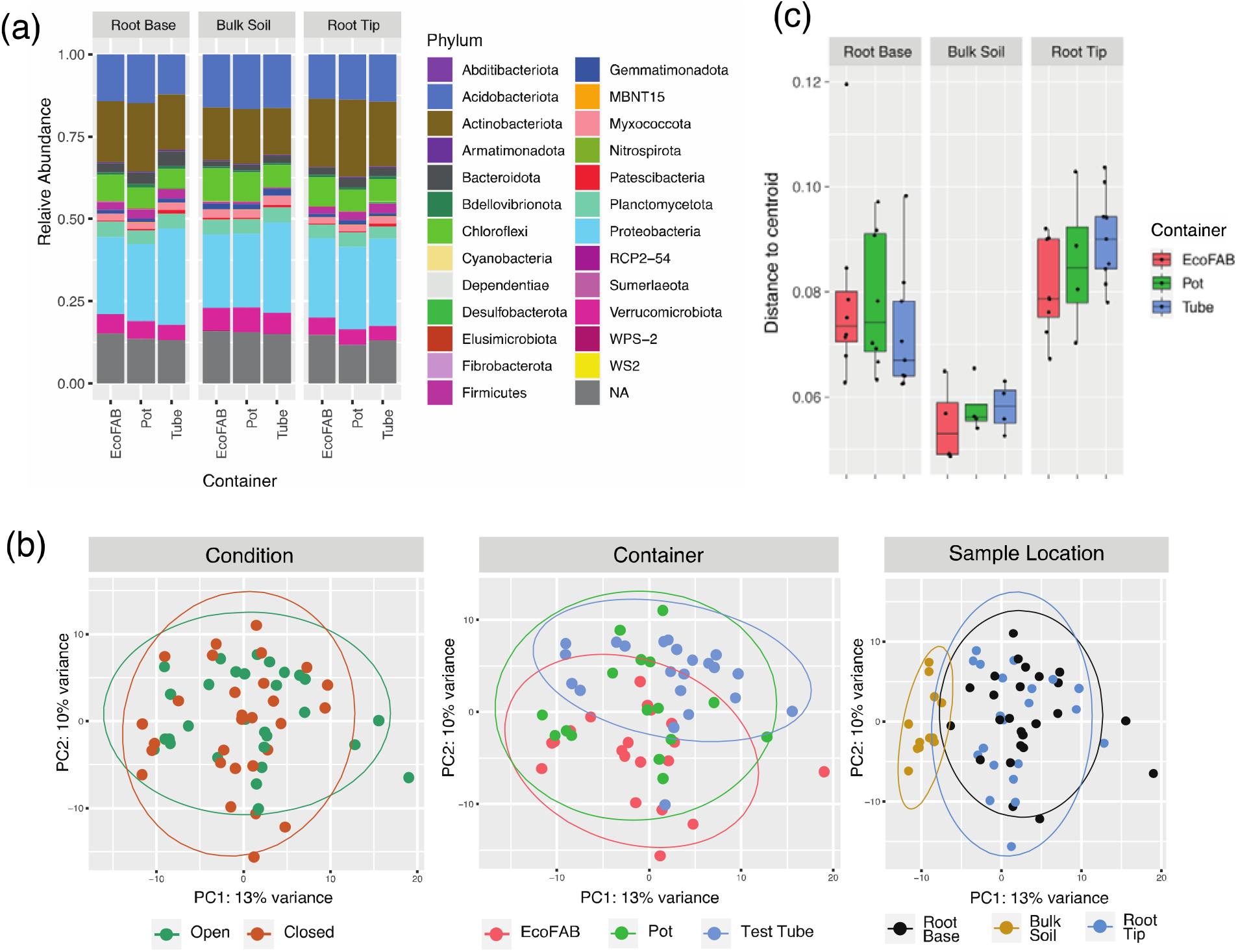
(a) Microbial relative abundance based on 16s rRNA amplicon sequencing of rhizosphere (root tip and root base) and bulk soil samples 14-day old *Brachypodium distachyon* grown in three different containers: EcoFAB, pot and test tubes, (b) PCA plot of variance stabilized 16S amplicon data, the samples are then visualized according to the three different variables examined: condition, container or sample location, (c) Boxplot showing multivariate homogeneity of group dispersions grouped according to sample location and container.

Comparative analysis of OTUs between different samples was then carried out to investigate the influence of three parameters tested, i.e., container type, location on root and open or closed condition. Principal Components Analysis (PCA) of the samples showed no clear separation among the two conditions or among the three container types whereas a distinct separation was observed between bulk soil samples compared to root base or root tip (**Fig. 1b**). However, no distinction was seen when comparing root base and root tip based on ordination analysis. This was supported statistically using MANOVA/*adonis* which showed the highest dissimilarity contributed by sample location (R^2^=0.10934, p=9.99e-05) followed by container type (R^2^=0.06336, p=0.00069) but no significant dissimilarity caused by either open or closed conditions (R^2^=0.02149, p=0.8119). Next, we examined whether the homogeneity within samples could be influenced by container type. Overall, the EcoFAB samples exhibited a comparable homogeneity among replicates of the same sample locations compared to the other two conventional containers such as pots and test tubes (**Fig. 1c**).

### 3.3 Pairwise comparison between sample locations showed the same differentially abundant OTUs regardless of container type

The OTUs which showed a statistically significant change in any of the pairwise comparisons, regardless of the containers, were selected and visualized using a neighbor-joining tree (**Fig. 2**). Distinct log-fold changes could be observed for comparisons looking at rhizosphere (root base or root tip) vs bulk soil. Further, analysis with Spearman’s correlation coefficient showed that the overall log-fold changes of each OTU were statistically positively correlated in most comparisons regardless of container (**Table S1**), with the only exception being the root tip vs root base changes observed in pot vs test tube (rho = -0.02, p = 0.78). In all three comparisons, results from EcoFAB samples were consistent with the others.

**Figure 2.**
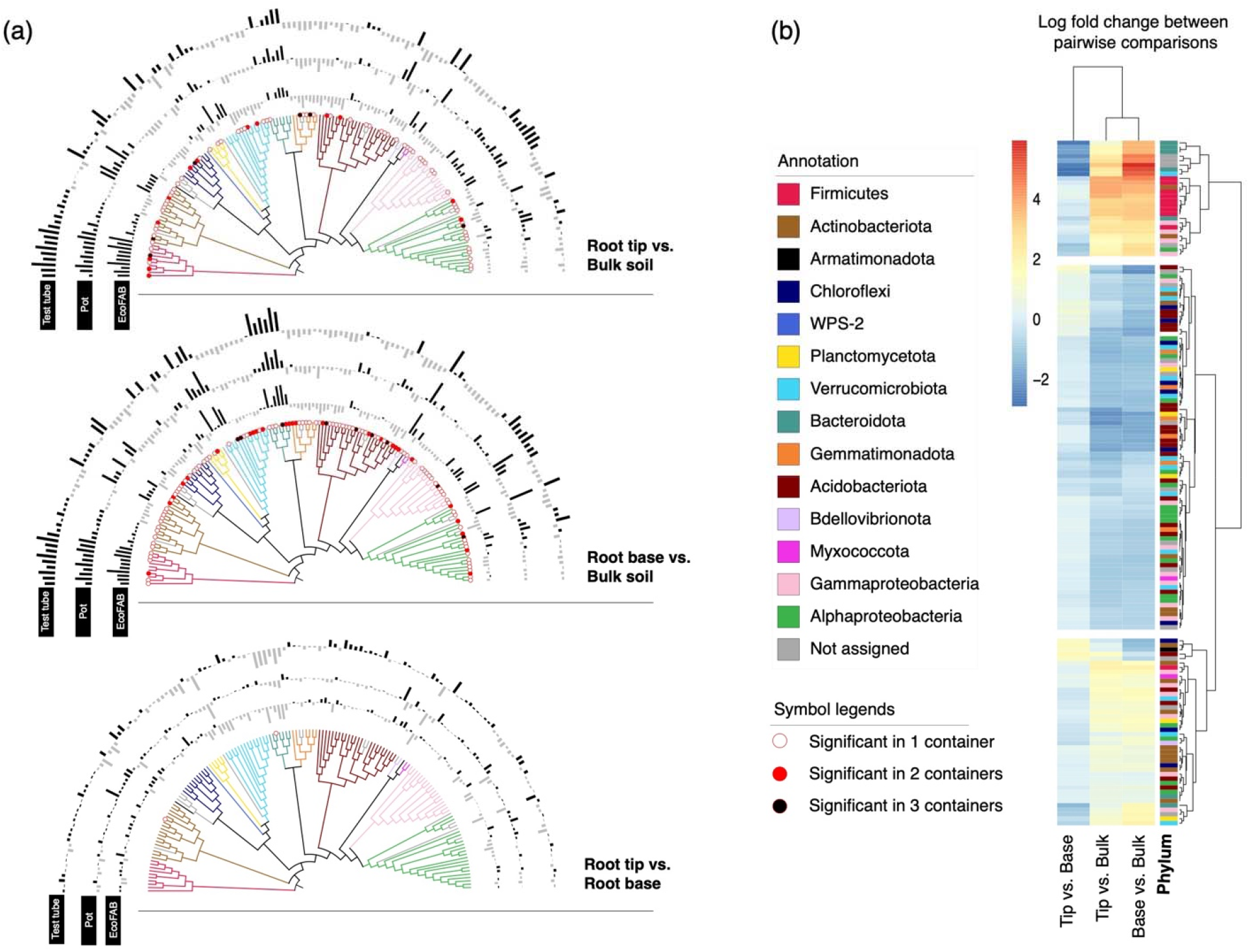
(a) Neighbor joining tree of selected OTUs which showed significant log fold changes during pairwise analysis of sample locations. The top tree depicts a pairwise comparison between root tip and root base and the bottom tree depicts the comparison between root base and bulk soil. The bar chart around the tree corresponds to log fold changes for each OTU in each of the different containers - test tube, pot or EcoFAB. An outward bar away from the tree represents a positive log fold change in the and an inward bar towards the tree represents a negative fold change in the respective OTU. The significant changes are indicated at the bottom of each node with a symbol. No symbol at the bottom of the node means the fold change is not statistically significant. (b) Clustering of selected OTUs based on pairwise comparison between sampling locations (ignoring containers) reveals three different clusters. Each OTU is colored by the phylum it belongs to.

Using comparisons solely based on sample location, the OTUs could be grouped into three distinct clusters (**Fig. 2b**). The first and smallest cluster showed the OTUs exhibiting significant increase in the rhizosphere (root base or root tip) compared to the bulk soil. Among them are multiple OTUs belonging to *Mucilaginibacter* (Bacteriodota), *Bacillus* (Firmicutes), *Paenibacillus* (Firmicutes), and unclassified Oxalobacteraceae (Gammaproteobacteria). The biggest cluster was for OTUs with a large decrease in the rhizosphere which included the phyla Acidobacteriota, Gemmatimonadota and Chloroflexi. The third cluster contained OTUs with minimal increase or decrease compared within sample locations and contained a mix of phyla.

### 3.4 Taxonomic analysis from metagenomics shows similar community composition to 16S rRNA based amplicon data

Read data from shotgun metagenome samples was directly assessed for bulk taxonomic composition using the ribosomal protein L6 (rpL6) marker gene. The phylum-level relative abundance in all samples showed dominance by the Proteobacteria, Actinobacteriota, Acidobacteriota, Planctomycetota and Verrucomicrobiota (**Fig. S3a**), similar to the 16S rRNA based community composition (**Fig. 1a**). A PCA plot also illustrated a clustering of the bulk soil samples distinctly from the rhizosphere samples as seen earlier in the corresponding 16S amplicon data (**Fig. S3b**). Overall, the metagenomic taxonomy was in correspondence with the 16S amplicon data and both types of analysis revealed minimal changes contributed by container differences.

### 3.5 Metagenome assembled genomes (MAGs) represent a small fraction of the total reads

Out of the 32 representative MAGs generated from 9 metagenomes after dereplication and quality filtering (**Fig. 3**), 11 MAGs belonged to Actinobacteriota; 6 MAGs from Gammaproteobacteria; 4 MAGs from Acidobacteriota and Alphaproteobacteriota; 3 MAGs each from Chloroflexota; 2 MAGs from Myxococcota and 1 MAG each from Gemmatimonadota and Elusimicrobiota (**Table S6**). As expected in systems with higher diversity, the total coverage of these genomes was rather low, representing ∼3% of the read data. 10 MAGs were identified to be differentially abundant across sample locations (**Fig. 3**). It is interesting to note that one Acidobacterial MAG (*Edaphobacter sp*.) had increased abundance in root tip compared to both bulk and base. Members of *Edaphobacter* genus are reported to be associated with ectomycorrhizal fungi and are important in their root colonization [41]. Only 6 MAGs had 16S rRNA and all these sequences had a 97-100% match with OTUs obtained from amplicon sequencing and similar phylogenetic classification.

**Figure 3.**
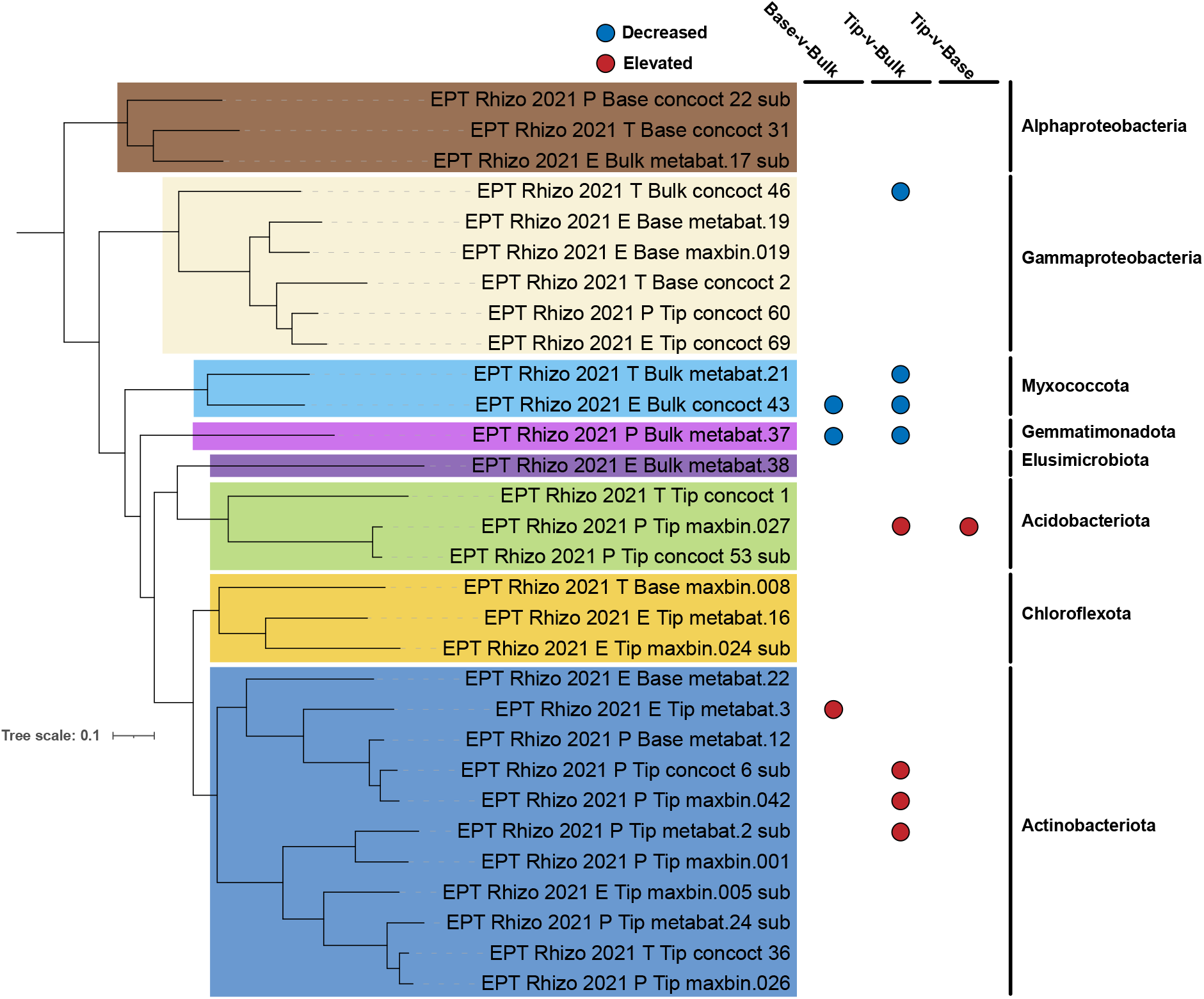
Phylogenetic tree of 30 of 32 dereplicated MAGs passing tree building criteria (*P*. calidifontis - GCA000015805 included as outgroup for rooting; not displayed) along with their differential abundance (significantly elevated or decreased; Wald Test - FDR ≤ 0.05) based on sample location. MAG names are colored based on their phylum-level classification and phyla names displayed on the right. Tree was inferred using a set of 25 phylogenetically informative marker genes conserved between Archaea and Bacteria.

### 3.6 Metagenome analysis reveals metabolic differences between root tip and bulk

5783 unique KEGG orthology groups (KOs) were annotated in the metagenomes, accounting for ∼30% of the total proteins predicted in each metagenome. PCA plot of KEGG Orthology (KO) composition of samples indicated that samples cluster by location irrespective of the container type (**Fig. S4**) and hence container parameter was excluded from further DESeq analysis. There were no differentially abundant KOs when root tip was compared to base, in congruence with observations from PCA analysis of OTUs (Section 3.2). Among the 55 differentially abundant KOs identified (**Fig. 4, Table S7**), 27 were enriched in root tip compared to bulk, while other 27 were decreased in tip vs. bulk and 2 KOs (one KO shared with decreased tip vs. bulk comparison) increased in bulk over base.

**Figure 4.**
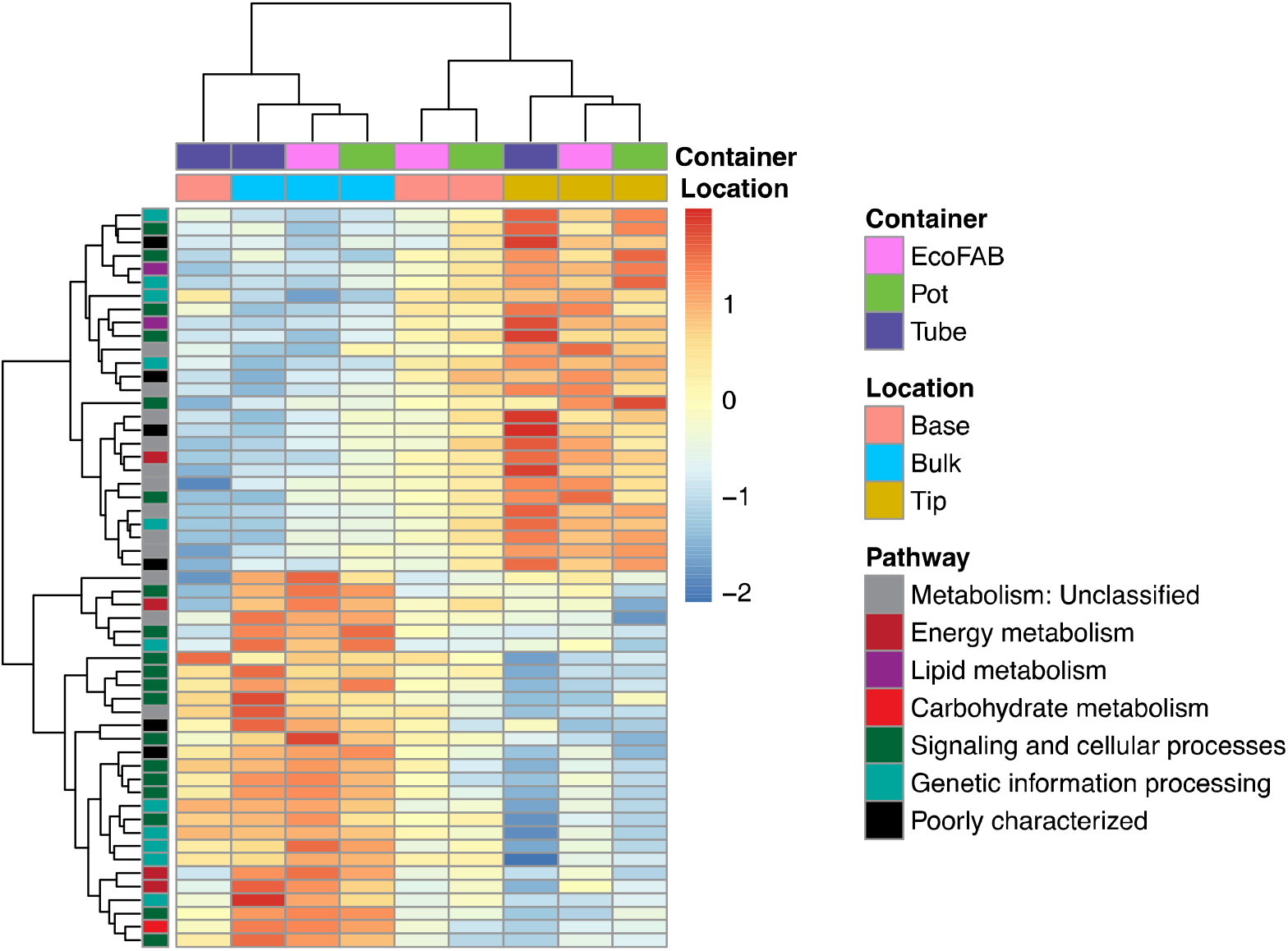
Heatmap of abundance of 55 differentially abundant KEGG Orthology genes across different locations in the 9 metagenome samples based on DESeq analysis (corrected p-value <0.1), normalized by z-score across all datasets. Each row represents a gene, colored by its KEGG level I classification. 27 KOs were enriched in root tip compared to bulk, 27 KOs were enriched in bulk compared to tip and 2 KOs in bulk over base.

KOs involved in different metabolic pathways were over-represented in tip compared to the bulk suggesting an active microbial population utilizing plant-derived compounds. These KOs, which could be broadly categorized as either enzymes, transcriptional regulators or transporters, play a critical role in substrate utilization as well as root colonization. Enzymes encoded were peptidases (*ampS, cwlO*), nucleases (*nucS*), kinases (*rsbW, fakA*), and other enzymes involved in fatty acid degradation (*acd*), lipid storage (*tgs/wax-dgat*), cell wall synthesis (*tagTUV*), and redox regulation (*gshA, fqr*). Transcriptional factors/regulator genes enriched in root tips were involved in regulation of purine catabolism (*pucR*), arabinogalactan biosynthesis (*embR*), biofilm formation (*sigB*) [42], sulfur utilization (sutR) and other functions (tetR). The enzyme, peptidoglycan DL-endopeptidase encoded by *cwlO*, has been shown to regulate biofilm formation and consequently root colonization in plant-beneficial rhizobacterium *Bacillus velezensis* SQR9 [43]. Interestingly, the anti-sigma factor *rsbW* and sigma factor *sigB* were identified as adjacent genes of *sigB* gene cluster and play important roles in stress resistance, biofilm formation and root colonization in *Bacillus cereus* 905 [42].Transporters involved in acquisition of copper (*ycnJ*), amino acid translocation (*rhtA*), ion transport (*nhaA*), and other nutrients (MFS (*mmr*) and ABC transporters (*mlaD/linD*)) were elevated in root tips. 4 other poorly characterized genes and gene involved in oxidative phosphorylation (*qcrC*) were also increased in the root tips over bulk metagenomes.

Microbes in the bulk soil do not have ready access to the labile carbon and nitrogen compounds in the exudate and hence may have to invest more in the biosynthesis of machinery for degradation of recalcitrant substrates and nutrient acquisition. Specifically, this involves KOs corresponding to transfer RNA biogenesis (*mnmE/trmE, gidA/mnmG*), transcriptional regulation (*rho, ada*), ribosome biogenesis (*rlmI*), and sulfur metabolism (*dmsBC*). Genes involved in heme uptake (*exbBD* and *tonB*) [44], nitrogen assimilation/quorum sensing (*rpoN*) [45] lipopolysaccharide export (*lptF*) were also increased in bulk soil. In addition, KOs involved in glycogen synthesis (*glgA*), polysaccharide biosynthesis/export (*wza/gfcE*), maintenance of cellular integrity under acidic stress (*ompA-ompF* porin), production of coenzymes (*pqqL*) involved in free-radical scavenging, regulation of exopolysaccharide production (*hprK*), periplasmic divalent cation tolerance (*cutA*) and osmotic stress genes (*osmY*) may confer resistance to environmental stressors like osmotic stress and desiccation [46, 47] present in bulk soil.

## 4. Discussion

We investigated the utility of EcoFABs as a possible alternative to conventional containers such as pots and tubes in studying the spatial microbial biogeography of the rhizosphere. Although, studies have shown that container design parameters such as size, density, depth can affect root growth and basic plant physiological traits during early developmental stages [5–9], our study in stark contrast, showed that EcoFABs had no significant impact on phenotypic plant growth. While most of these studies looked at container sizes around 50cm^3^, these studies were performed using woody tree seedlings such as *Pinus sp*. (Pine tree species) and *Quercus sp*. (Oak tree species). Container impacts may not apply to softer wheat plants such as *B. distachyon* to a discernible extent. This emphasizes the importance of using the correct standardized containers to perform accurate study comparisons for the system under investigation.

Next, we investigated the impact of microbial community assembly on the root impacted by container differences using both 16S amplicon sequencing and metagenomics. Based on 16S amplicon sequencing results, the microbial community of each location with respect to root showed relatively similar composition across all containers. Differences were observed mostly in root tip or base locations compared to the bulk soil. At root tips, a decrease of bacterial OTU richness and alpha diversity when compared to bulk soil has been previously reported [3, 48]. This reduction in microbial diversity in the rhizosphere is commonly observed [49] as the root exudates create a selective environment, recruiting selected microbes from bulk soil. We further observed that even within the rhizosphere, root tips had lower bacterial diversity (richness and eveness) than root base, which concurs with the other studies conducted on *Brachypodium* roots [50, 51]. Root tip environment appears to be more stochastic compared to the root base as the assembly patterns appear to be more deterministic in older parts of the root [49]. This is true in our study as well, there were a higher number of significant OTUs in the comparison of base vs bulk than comparing tip vs bulk (Fig. 2a). Nonetheless, overall correlations show a significantly positive correlation which meant that the rhizosphere effect is already developing at the tip even for 2 week old seedlings of *Brachypodium*. Usually, microbial composition studies tend to occur at later stages of *Brachypodium* growth [50–52] because the plant often takes 30 – 35 days to reach maturity [40]. Our study, however, shows that a rhizosphere effect may be occurring as early as 14 days into the plant growth albeit a weaker impact at the root tips.

Only some of the dominant rhizosphere community members such as Gammaproteobacteria and Bacteriodota matched the observations in a previous study with *Brachypodium* rhizosphere [50]. Phyla such as *Betaproteobacteria*, which were highly enriched in a previous study with mature plants [50], were neither abundant nor showed enrichment in the rhizosphere. Nonetheless, other rhizosphere enriched groups in this study include Actinobacteria, Acidobacteria and Verrucomicrobia which seems to be more of an effect of the low pH soil characteristic of our field site [53]. Additionally, in that study [50] *Brachypodium* was grown in sand amended soil which could explain the differences. Actinobacteria, for instance, is associated with rhizosphere in soils with high organic content [54, 55]. In another study where fine scale sampling of 4-week-old *Brachypodium* roots was performed, Firmicutes were more abundant in root tips compared to root base, whereas opposite trend was observed for Verrucomicrobia [51]. Phyla such as Actinobacteria, Proteobacteria and Bacteriodota were reported to be enriched in wheat rhizosphere [56]. Thus, in line with prior studies, our data also suggests that a combination of root exudates and edaphic factors are working in tandem to enrich a specific rhizosphere community.

Among 150 OTUs which were differentially abundant between different sampling locations, all OTUs belonging to phylum Firmicutes and Bacteriodota were enriched in rhizosphere over bulk soil. These included genera *Bacillus* and *Paenibacillus* (Firmicutes) and *Mucilaginibacter* (Bacteriodota). Members of *Paenibacillus* have been isolated from rhizosphere of wide variety of plants; several of these are capable of fixing-nitrogen [57–59]. Similarly, several *Mucilaginibacter* strains have been isolated from rhizosphere, and a comparative analysis of various strains in this genus highlighted the presence of diverse carbohydrate active enzymes including cellulose-degrading enzymes [60]. Impacts of different Bacillus isolates on *Brachypodium* plants have been characterized previously; *Bacillus* isolates can accelerate growth, provide drought protection [61], influence root architecture [61] and can modulate plant hormone homeostasis. Some *Bacillus*, could be classified as r-strategists, which can quickly grow in response to nutrient availability in rhizosphere [51].

Majority of differentially abundant OTUs belonging to Gemmatimonodota, Acidobacteria and Verrucomicrobia had reduced abundance in the rhizosphere compared to bulk. These bacterial groups are slow-growing and oligotrophic [62–64], thus more suited to survive in bulk soil away from the nutrient-rich rhizosphere. On the contrary, the OTUs belonging to Actinobacteria, Gammaproteobacteria and Alphaproteobacteria showed no clear trends—OTUs could be either enriched or depleted in the rhizosphere.

We observed congruence between taxonomic results obtained by 16S rRNA gene sequencing and metagenomics (rpL6 marker gene), demonstrating reliability of different sequencing methodologies for bacterial profiling (short read Illumina vs. long-read technology). Comparative analysis of metagenomic functional potential between various sampling locations revealed significant differences between root tips and bulk soil. KO genes involved in different metabolic pathways and root colonization were over-represented in tip compared to bulk suggesting an active microbial population capable of utilizing plant-derived exudates and occupying the rhizosphere. KO genes associated with biosynthesis of machinery for degradation of recalcitrant substrates, nutrient acquisition and stress-tolerance were prominent in bulk soil where readily available substrates are scarce in comparison to the vicinity of roots. These findings are consistent with other metagenomic studies comparing rhizosphere vs. bulk soil [65] and also in agreement with the results from 16S amplicon sequencing, where rhizosphere is abundant in fast-growing groups and bacterial assembly in root tips is stochastic, while bulk soil is enriched with groups that are more oligotrophic and adapted to survive in nutrient-limited conditions.

We would also like to highlight a few shortcomings of this study. As a result of low DNA yields, samples were pooled for metagenomics which led to low sample numbers. In addition to this, genome-resolved metagenomics yielded fewer genomes making statistical analysis of genome relative abundance and metabolic enrichment analysis difficult. Differences in gene abundance were observed only between root tips and bulk soil, thus differences within rhizosphere compartments (tip vs. base) are unclear which is probably due to sampling of young plants. This is inturn associated with EcoFAB size which limits how long plants could be grown, but can be easily addressed with bigger molds.

Thus, we have demonstrated the influence of root exudation patterns in shaping microbial communities on different sections of the root in comparison with bulk soil in as young as 14-day old *Brachypodium* plants through 16S rRNA amplicon sequencing and metagenomic analyses. To further probe into the physiology of root-enriched microbes, we will perform high-throughput enrichment of this rhizobiome on known root exudate compounds to create reduced complexity communities. This biogeography study serves as proof of concept for further investigation into high-resolution sampling of rhizosphere to understand biological interactions occurring at finer scales. We are currently working on engineering materials that can be integrated into EcoFABs to enable localized, sub-millimeter scale sampling at different timepoints.

## 5. Acknowledgements

This research work was funded through the Microbial Community Analysis and Functional Evaluation in Soils (m-CAFEs) Science Focus Area Program at Lawrence Berkeley National Laboratory funded by the U.S. Department of Energy, Office of Science, Office of Biological & Environmental Research Awards DE-AC02-05CH11231.

## 7. Data Availability Statement

The 16S rRNA amplicon sequences and metagenome-assembled genomes generated during the current study are available in the NCBI SRA repository, under the BioProject ID PRJNA902408. The full assemblies for each metagenome sample are publicly available at our in-house analysis platform, ggKbase (https://ggkbase.berkeley.edu).

