## Supplementary material 1 for "Fine scale sampling reveals spatial heterogeneity of rhizosphere microbiome in young *Brachypodium* plants"

**Materials and methods: *2.7 Phylogenetic and abundance analysis of genome bins***

Phylum level taxonomic assignments of de-replicated genome bins were inferred using GTDB-Tk v1.5.1 [[1]](https://sciwheel.com/work/citation?ids=7833838&pre=&suf=&sa=0&dbf=0) with reference data version r202 and the following options: classify_wf --genome_dir [input genomes] --out_dir [output folder] -x fa --cpus 20. Phylogenetic relationships between de-replicated genome bins were inferred using GToTree v1.5.22 using the following parameters: -H Bacteria_and_Archaea.hmm -j 8 -d. Briefly, a set of 25 phylogenetically informative marker genes indicated in the GToTree Bacteria_and_Archaea marker set were first identified in our 32 de-replicated genomes and 1 genome (*P.* calidifontis - GCA000015805) included as an outgroup for analysis. To be included in phylogenetic analysis ≥ 50 % of marker genes needed to be identified, two genomes in our analysis did not pass this criterion and were omitted from the final phylogenetic tree. Marker gene sequences were independently aligned, and a phylogenetic tree was produced using FastTree2 [[2]](https://sciwheel.com/work/citation?ids=178753&pre=&suf=&sa=0&dbf=0). The tree was displayed and rooted in Geneious Prime v2020.2.4. For display in the manuscript, the outgroup organism was removed and phylum level lineages were colored based on GTDB-Tk taxonomy assignment with the exception of one MAG (EPT_Rhizo_2021_E_Tip_metabat.16) where tree-based taxonomy indicated clustering within the Chloroflexota rather than the GTDB-Tk inferred phylum of Actinobacteriota.

The relative abundance of the 32 species level genome bins in all samples was assessed by cross mapping reads from each of the 9 samples back to the genome bins using bowtie2 with default options. Read counts and coverage of genomes in each sample were quantified from bowtie2 outputs using coverM (https://github.com/wwood/CoverM) in genome mode and requiring ≥ 95 % read mapping identity. Differential abundance of genomes between rhizosphere spatial locations was assessed using the DESeq2 package in R [[3]](https://sciwheel.com/work/citation?ids=129353&pre=&suf=&sa=0&dbf=0). Briefly, counts of mapped reads per genome per sample (n = 9 samples) were imported into R and analyzed with a DESeq2 design formula of the form: counts ~ container type + location. This design assesses variance in counts based on location while controlling for count variance introduced by the three container types. Pairwise comparisons between the three different possible location parings were carried out using the DESeq2 results function with contrast argument specifying the pairings. Differential abundance was considered significant when the FDR corrected p-value for a genome in a contrast was ≤ 0.05.

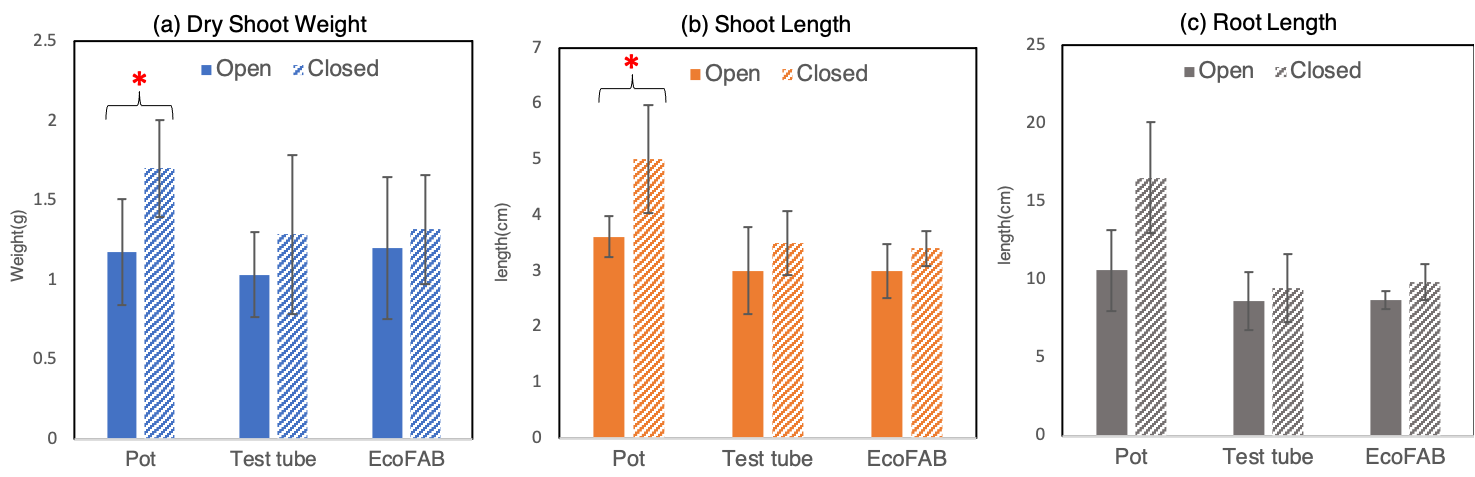

**Figure S1**. Phenotypic characteristics, namely (a) dry shoot weight, (b) shoot length and (c) root length of 14-day old *Brachypodium distachyon* grown in three different containers; pot, test tube and EcoFAB.

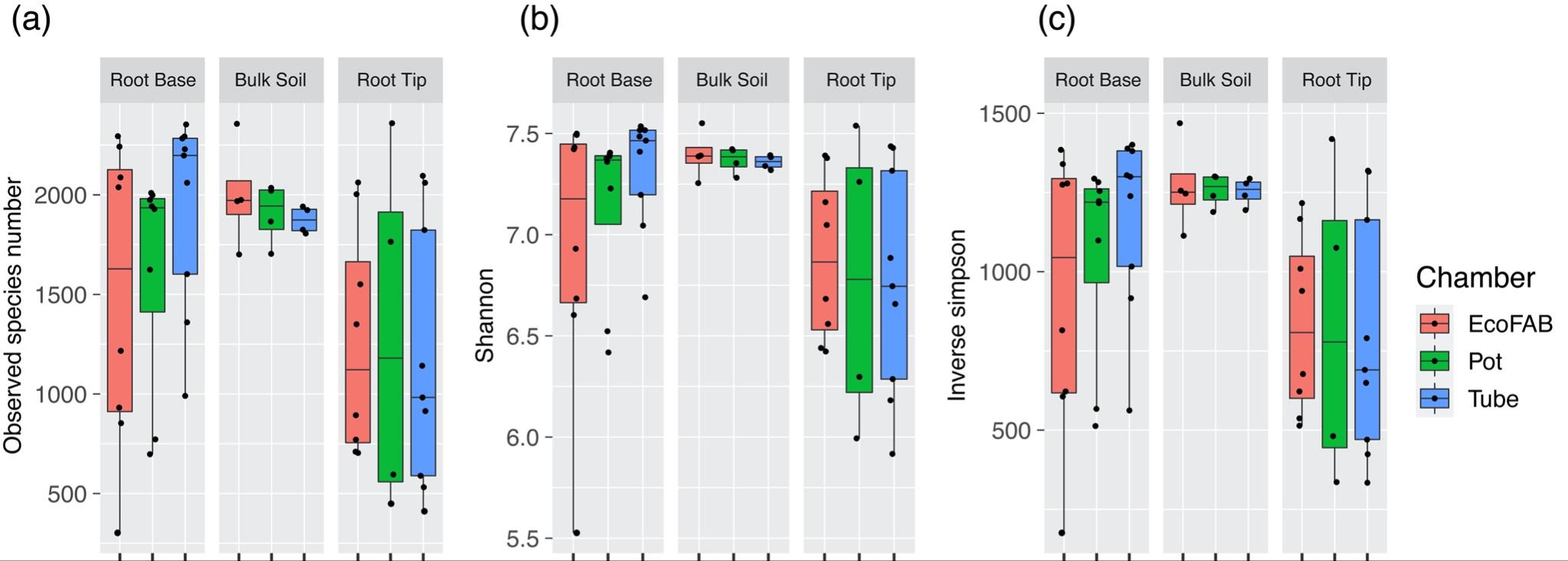

**Figure S2**. Three different Alpha diversity indices, (a) observed species number, (b) Shannon and (c) Inverse Simpson indices of the microbial community on the rhizosphere and bulk soil of *B. distachyon* grown in three different containers/chambers.

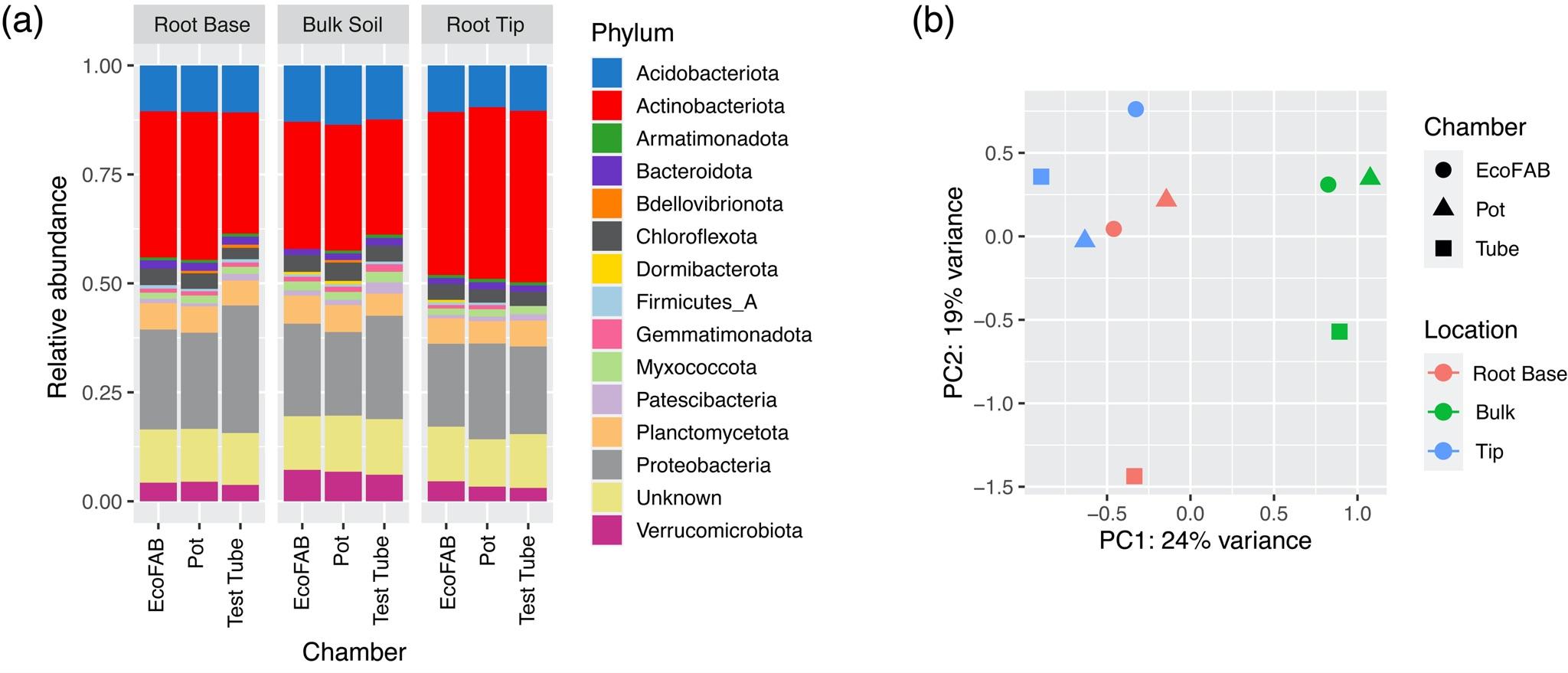

Figure S3. (a) Microbial taxonomic relative abundance based on RPL6 gene from metagenomics data of rhizosphere and bulk soil samples of *B. distachyon*; only the top 15 abundant phyla are shown here; (b) PCA plot of variance transformed RPL6 gene based taxonomic data.

Table S1. Pairwise comparison of log-fold changes of 150 OTUs between different locations and container types using Spearman’s correlation

| Location | Container | Rho value | p-value |
| --- | --- | --- | --- |
| Root base vs. Bulk soil | EcoFab vs. Test Tube | 0.759893 | 1.77E-29 |
|  | EcoFab vs.Pot | 0.78196 | 3.46E-32 |
|  | Test Tube vs.Pot | 0.749973 | 2.37E-28 |
| Root tip vs. Bulk soil | EcoFab vs. Test Tube | 0.759303 | 2.08E-29 |
|  | EcoFab vs.Pot | 0.692124 | 1.04E-22 |
|  | Test Tube vs.Pot | 0.696885 | 4.01E-23 |
| Root tip vs. Root base | EcoFab vs. Test Tube | 0.177475 | 0.029803 |
|  | EcoFab vs.Pot | 0.184344 | 0.02393 |
|  | Test Tube vs.Pot | -0.02207 | 0.788657 |

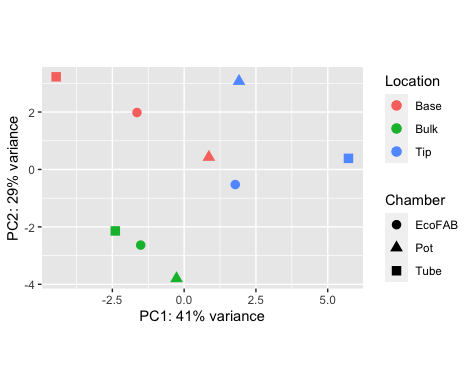

Figure S4. PCA plot of variance transformed KEGG Orthology (KO) gene composition of samples.
